# Effect of task and attention on neural tracking of speech

**DOI:** 10.1101/568204

**Authors:** Jonas Vanthornhout, Lien Decruy, Tom Francart

## Abstract

EEG-based measures of neural tracking of natural running speech are becoming increasingly popular to investigate neural processing of speech and have applications in audiology. When the stimulus is a single speaker, it is usually assumed that the listener actively attends to and understands the stimulus. However, as the level of attention of the listener is inherently variable, we investigated how it affected neural envelope tracking. Using a movie as a distractor, we varied the level of attention while we estimated neural envelope tracking. We varied the intelligibility level by adding stationary noise. We found a significant difference in neural envelope tracking between the condition with maximal attention and the movie condition. This difference was most pronounced in the right-frontal region of the brain. The degree of neural envelope tracking was highly correlated with the stimulus signal-to-noise ratio, even in the movie condition. This could be due to residual neural resources to passively attend to the stimulus. When envelope tracking is used to objectively measure speech understanding this means that the procedure can be made more enjoyable and feasible by letting participants watch a movie during stimulus presentation.

## 1 Introduction

EEG-based measures of neural tracking of natural running speech are becoming increasingly popular to investigate neural processing of speech and for applications in domains such as audiology (e.g. Vanthornhout et al. (2018); Lesenfants et al. (2019)) and disorders of consciousness (e.g. Braiman et al. (2018)). With the stimulus reconstruction method, natural running speech is presented to a listener and a feature of the speech signal, such as the envelope, is reconstructed from the brain responses. The correlation between the original and reconstructed signal is a measure of neural coding of speech (Ding and Simon, 2012). Inversely, the brain responses can be predicted from features of the speech signal. Lalor et al. (2009), for example, found that the transfer function from the acoustic envelope to brain responses, also called the temporal response function (TRF), exhibits interpretable peaks, such as the N1 and P2 peaks (cf. infra).

These methods, reconstructing the stimulus or predicting the brain responses, are often used with either a single speaker or in a competing talker paradigm. With a single speaker, it is usually assumed that the listener actively attends to and understands the stimulus. In the competing talker paradigm, the listener’s task is usually to attend to one of several speakers. However, in either paradigm, it is possible that a listener tunes out once in a while. In this study, we investigated the effect of attention on neural tracking, in function of the signal-to-noise ratio. Participants paid full attention or we distracted them by letting them watch a movie. We did this for two types of stimuli, a story and repeated sentences.

Attention is a conscious active process and has a strong top-down effect on processes along most of the auditory pathway (Forte et al., 2017; Fritz et al., 2007). The concept of auditory attention can be illustrated by the cocktail party problem described by Cherry (1953). Although it is possible to listen to a certain speaker while ignoring other speakers, some important information, such as your name, can still be recognised when uttered by one of the ignored speakers. The combination of bottom-up salience and top-down attention leads to speech understanding (Shinn-Cunningham, 2008).

The effect of attention has already been investigated in objective measures of hearing, such as auditory brainstem repsonses (ABRs), auditory steady-state responses (ASSRs) and late auditory evoked potentials. It is often assumed that attention has a minimal effect on the ABR, as it can even be measured when the subject is asleep. However, the results are inconclusive for short clicks, pure tones and speech (Brix, 1984; Hoormann et al., 2000; Galbraith et al., 2003; Varghese et al., 2015; Lehmann and Schönwiesner, 2014).

The effect of attention has also been investigated for ASSRs. Ross et al. (2004) used a modulation discrimination task to focus auditory attention on modulated tone stimuli. They found a significant effect of focused auditory attention on the 40-Hz ASSR amplitude. Roth et al. (2013b) measured 40-Hz ASSRs, while their participants played a computer game (Tetris), and investigated the effect of game difficulty level on the SNR of the 40-Hz ASSR. They found a significant reduction in SNR with increase in difficulty. This is consistent with the theory that irrelevant stimuli are suppressed when neural resources are required for another task (Hein et al., 2007). Müller et al. (2009) found some differences at 20 Hz. However, other studies found no effect of attention: Skosnik et al. (2007) did not find a difference at 20 Hz, Linden et al. (1987) did not find a difference at 40 Hz. A reason can be that the participants did not do what was expected from them. Furthermore, these differences could be due to differences in the attentional tasks used. Overall, there is no consensus on what the effect is of attention on neural tracking of the stimulus.

Late auditory evoked potentials allow investigation of the time-domain brain response to repeated transient sounds. As attention originates in the cortex (Fritz et al., 2007), we expect that mainly later responses are modulated by attention as they have a cortical origin. The N1 and P2 components have been shown to be enhanced by attention to the stimulus (Astheimer and Sanders, 2009; Picton and Hillyard, 1974; Näätänen et al., 2011), and are also influenced by the visual load (Regenbogen et al., 2012).

As the human brain is finely attuned to speech perception, it is interesting to investigate the effect of attention on neural measures of speech tracking, which can be used with natural speech as the stimulus, in contrast to the artificial repeated stimuli used in the paradigms described above. There is a wide range of data available on neural tracking of speech in a competing talker paradigm. Two speakers or more are presented simultaneously to a listener, whose task it is to attend to one of them. A stimulus feature is then decoded from the EEG and correlated with the features of each speaker. A decoder transforms the multi-channel neural signal into a single-channel feature, by linearly combining amplitude samples across sensors and across a post-stimulus temporal integration window. Comparison of these correlations yields information on which speaker was attended. It has been shown that the attended speaker can be decoded with high accuracy (in the order of 80-90% for trials of 20-30s) (e.g. O’Sullivan et al. (2015); Mirkovic et al. (2015); Das et al. (2016, 2018)).

By changing the number of EEG samples included in the decoder, i.e. the latencies of EEG respective to the stimulus, it can be investigated at which time lags (and therefore neural sources) the effect of attention appears. Not unexpectedly, attention mostly has an effect at longer latencies, ranging from 70 ms to 400 ms (Hillyard et al., 1973). Ding and Simon (2012) investigated the neural responses in dual-talker scenario and found that peaks in the TRF at 100 ms were more influenced by attention than peaks at 50 ms. Other experiments show effects at a later stage, beyond 150 ms (Snyder et al., 2006; Ross et al., 2009). O’Sullivan et al. (2015) found best attention decoding performance with latencies around 180 ms. Moreover, Puvvada and Simon (2017) found that early responses (< 75 ms) are minimally affected by attention. However, even at the brainstem level, around 5 ms latency, modulation by attention can already be found, showing the top-down effects (Forte et al., 2017). As a conclusion, attention prominently affects the later responses but can also effect very early responses.

In a competing talker paradigm, it is also interesting to investigate the relative level of neural speech tracking: is the attended speaker represented more strongly, the unattended one suppressed, or a combination? Bidet-Caulet et al. (2010) shows that the representation of the attended stimulus is not enhanced, but responses to the unattended stimulus are attenuated. Irrelevant stimuli are actively suppressed. Melara et al. (2002) on the other hand show that excitatory (enhancing the attended stimulus) and inhibitory (attenuating the unattended stimulus) processes work interactively to maintain sensitivity to environmental change, without being disrupted by irrelevant events. Many researchers have found enhancement of the cortical envelope tracking of attended sounds, relative to unattended sounds. This attentional effect can be found as early as 100 ms post stimulus (Melara et al., 2002; Choi et al., 2014), corresponding with lags associated with the auditory cortex. Kong et al. (2014) however demonstrate that top-down attention can both enhance the neural response to the attended sound stream and attenuate the neural responses to an unwanted sound stream.

The literature on the effect of attention on neural tracking of a single speaker is more sparse. Kong et al. (2014) measured neural tracking of a single talker in quiet, while the participant either actively listened to a stimulus or watched a movie and ignored the auditory stimulus. They found that neural tracking, measured as the peak cross-correlation between EEG signal and stimulus envelope, was not significantly affected by listening condition. However, the shape of the cross-correlation function showed stronger N1 and P2 responses in the active listening condition and weaker P1 response compared to the movie condition. However, they did not include more intensive distractors, nor did they investigate the effect of speech intelligibility. Moreover, their analysis was limited to a cross-correlation while decoding the envelope from the neural responses would be more powerful as it uses information from all channels.

We investigated the effect of attention on neural envelope tracking. Attention was manipulated by letting the participants actively attend to the stimulus and answer comprehension questions, or watch a silent movie and ignore the acoustic stimulus. This was done at multiple levels of speech understanding, by adding stationary background noise.

We hypothesised that envelope tracking would be strongest in the attended condition. However, watching a movie may stabilize the level of attention, as maintaining focus on the movie might be easier than maintaining focus on (uninteresting) sentences, therefore reducing intra-subject variability. We also hypothesized that this effect would be most apparent for decoders with a temporal integration window beyond 100 ms.

## 2 Methods

### 2.1 Participants

We recruited 20 young normal-hearing participants between 18 and 35 years old. Every subject reported normal hearing (thresholds lower than 20 dB HL for all audiometric octave frequencies), which was verified by pure tone audiometry. They had Dutch (Flemish) as their mother tongue and were unpaid volunteers. Before each experiment the subjects signed an informed consent form approved by the local Medical Ethics Committee (reference no. S57102).

### 2.2 Apparatus

The experiments were conducted using APEX 3 (Francart et al., 2008), an RME Multiface II sound card (RME, Haimhausen, Germany) and Etymotic ER-3A insert phones (Etymotic Research, Inc., Illinois, USA) which were electromagnetically shielded using CFL2 boxes from Perancea Ltd. (London, United Kingdom). The speech was always presented at 60 dBA and the setup was calibrated with a 2 *cm*^3^ coupler (Brüel & Kjær 4152, Nærum, Denmark) using the speech weighted noise of the corresponding speech material. The experiments took place in an electromagnetically shielded and soundproofed room. To measure EEG we used an ActiveTwo EEG setup with 64 electrodes from BioSemi (Amsterdam, Netherlands).

### 2.3 Behavioural experiments

Behavioural speech understanding was measured using the Flemish Matrix test (Luts et al., 2015). Each Matrix sentence consists of 5 words spoken by a female speaker and was presented to the right ear. Right ear stimulation was chosen as this is how the Matrix test has been standardised. The sentences of the Flemish Matrix test have a fixed structure of ‘name verb numeral adjective object’, e.g., ‘Sofie ziet zes grijze pennen’ (‘Sophie sees six grey pens’). Each category of words has 10 possibilities. The result of the Flemish Matrix test is a word score.

Several lists of sentences were presented at a fixed SNR ranging from −12 dB SNR to 1 dB SNR (and for some participants (n = 8) also in silence). For the constant procedure, we estimated the speech reception threshold (SRT) of the participants by fitting a psychometric curve through their word scores, using the formula 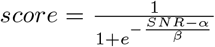, with *α* the SRT and *β* the slope. For some participants (n = 10) we used 3 runs of an adaptive procedure (Brand and Kollmeier, 2002) in which we changed the SNR until we obtained 29%, 50% or 71% speech understanding. We considered the SNR after the last trial as the SRT at the desired level.

### 2.4 EEG experiments

After the behavioural experiment, we conducted the EEG experiment. The stimuli were presented with the same set-up as the behavioural experiments, with the exception that we used diotic stimulation for the EEG experiment to make the experiments more comfortable.

#### 2.4.1 Speech material

We presented stimuli created by concatenating two lists of Flemish Matrix sentences (2 s per sentence) with a random gap of minimum 0.8 s and maximum 1.2 s between the sentences. The total duration of this stimulus was around 120 seconds with 80 seconds of speech. The stimulus was presented at different SNRs (−12.5 dB SNR up to +2.5 dB SNR and without noise). Each stimulus was presented 3 times. The total duration of the experiment was 2 hours excluding breaks. To keep the subjects attentive, they could take a break when desired. Also to keep the participants attentive, we asked questions during the trials without movie about the stimuli before and after the presentation of the stimulus. The questions were typically counting tasks, e.g. ‘How many times did you hear “red balls”?’. The answers were noted but were not used for further processing.

The participants also listened to the children’s story ‘Milan’, written and narrated in Flemish by Stijn Vranken (Flemish male speaker). It was approximately 15 minutes long and was presented without any noise. The purpose was to have a continuous, attended stimulus to train the linear decoder (cf. infra).

#### 2.4.2 Attention conditions

While listening to the Matrix sentences, the participants’ attention was manipulated using two different tasks. The participants were either instructed to (1) attentively listen to the sentences, and respond to questions and (2) watch a silent movie. We instructed the participants not to listen to the auditory stimulus when watching the movie.

In the movie condition, we used a subtitled cartoon movie of choice by the participant to aim for a similar level of distraction across subjects. The text dialogue effectively captures attention while not interfering with auditory processing. A cartoon movie was chosen to avoid realistic lip movements, which may activate auditory brain areas (O’Sullivan et al., 2017). Navarra (2003) investigated if incongruent linguistic lip movements interfere with the perception of auditory sentences, and found that visual linguistic stimuli produced a greater interference than non-linguistic stimuli. For some subjects *n* = 9, we also presented a movie during a story stimulus. This story stimulus was a 15 minutes excerpt from ‘De Wilde Zwanen’, however it was different from the previously mentioned story stimulus.

### 2.5 Signal processing

All signal processing was implemented in MATLAB R2016b (The MathWorks, Inc.).

#### 2.5.1 Speech

We measured neural envelope tracking by calculating the correlation between the stimulus speech envelope and the envelope reconstructed using a linear decoder.

The speech envelope was extracted from the stimulus according to Biesmans et al. (2017), who investigated the effect of envelope extraction method on auditory attention detection and found best performance for a gammatone filterbank followed by a power law. In more detail, we used a gammatone filterbank (Søndergaard and Majdak, 2013; Søndergaard et al., 2012) with 28 channels spaced by 1 equivalent rectangular bandwidth, with center frequencies from 50 Hz to 5000 Hz. From each subband we extracted the envelope by taking the absolute value of each sample and raising it to the power of 0.6. The resulting 28 subband envelopes were averaged to obtain one single envelope.

To remove glitches, we blanked each sample having a higher amplitude than 500 *μ*V. These blanked portions were linearly interpolated. To decrease processing time the EEG data and the envelope were downsampled to 256 Hz. To reduce the influence of ocular artefacts, we used a multi-channel Wiener filter (Somers et al., 2018). This spatial filter estimates the artefacts, and subtracts them from the EEG. After artefact suppression, the EEG data were re-referenced to the average of the 64 channels.

The speech envelope and EEG signal were band-pass filtered. We investigated the delta band (0.5 - 4 Hz). The same filter (a zero phase Chebyshev type 2 filter with 80 dB attenuation at 10% outside the passband) was applied to the EEG and speech envelope. After filtering, the data were further downsampled to 128 Hz.

The decoder linearly combines EEG electrode signals and their time shifted versions to optimally reconstruct the speech envelope. In the training phase, the weights to be applied to each signal in this linear combination are determined. The decoder was calculated using the mTRF toolbox (version 1.1) (Lalor et al., 2006, 2009) and applied as follows. As the stimulus evokes neural responses at different delays along the auditory pathway, we define a matrix *R* containing the shifted neural responses of each channel. With *g* the linear decoder and *R* the shifted neural data, the reconstruction of the speech envelope 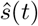 was calculated as

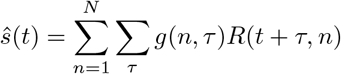

with *t* the time ranging from 0 to *T*, *n* the index of the recording electrode and *τ* the post-stimulus integration window length used to reconstruct the envelope. The matrix *g* can be determined by minimizing a least-squares objective function

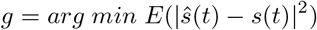

with *E* the expected value, *s*(*t*) the real speech envelope and 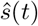 the reconstructed envelope. In practice we calculated the decoder by solving

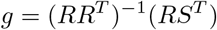

with *S* a vector of stimulus envelope samples. The decoder was calculated using ridge regression on the inverse autocorrelation matrix. We used post-stimulus lags of 0 - 500 ms.

We trained a new decoder for each subject on the story stimulus, which was 15 minutes long. After training, the decoder was applied on the EEG responses to the Flemish Matrix material. To measure the correspondence between the speech envelope and its reconstruction, we calculated the bootstrapped Spearman correlation between the real and reconstructed envelope. Bootstrapping was applied by Monte Carlo sampling of the two envelopes.

We also estimated the temporal response function on the stories and on the Matrix sentences in the speech in quiet condition. This is similar to the calculation of the decoder but we now predict EEG instead of reconstructing the envelope.

## 3 Results

### 3.1 Behavioural speech understanding

For each subject, we fitted a psychometric curve on the SNR versus percentage correct score and estimated the corresponding SRT and slope. The average SRT is −8.03 dB SNR (standard deviation: 1.27 dB) with an average slope of 14.1 %/dB (standard deviation: 3.4 %/dB).

### 3.2 Neural responses to speech

#### 3.2.1 Effect of SNR and attention

Figures 1a and 1b show neural envelope tracking as function of SNR using respectively a 0 - 75 ms and a 0 - 500 ms integration window in the delta band (0.5 - 4 Hz) for the 2 conditions: maximal attention and watching a movie. As we tested a wide variety of SNRs, we clustered the SNRs using k-means clustering in 7 clusters^1^. When multiple correlations per SNR were present for a subject, the average was taken. Visual assessment shows a systematic increase in neural tracking with SNR (related to intelligibility). Indeed, the Spearman correlations between SNR and neural envelope tracking for the attention and movie condition using a 0 - 75 ms integration window in the decoder are respectively: 0.38 (*p* < 0.001) and 0.52 (*p* < 0.001). These correlations are significantly different from each other *p* = 0.02 (Hittner et al., 2003; Diedenhofen and Musch, 2015). Using a 0 - 500 ms integration window they are respectively: 0.36 (*p* < 0.001), 0.48 (*p* < 0.001) and also significantly different *p* = 0.04.

**Figure 1:**
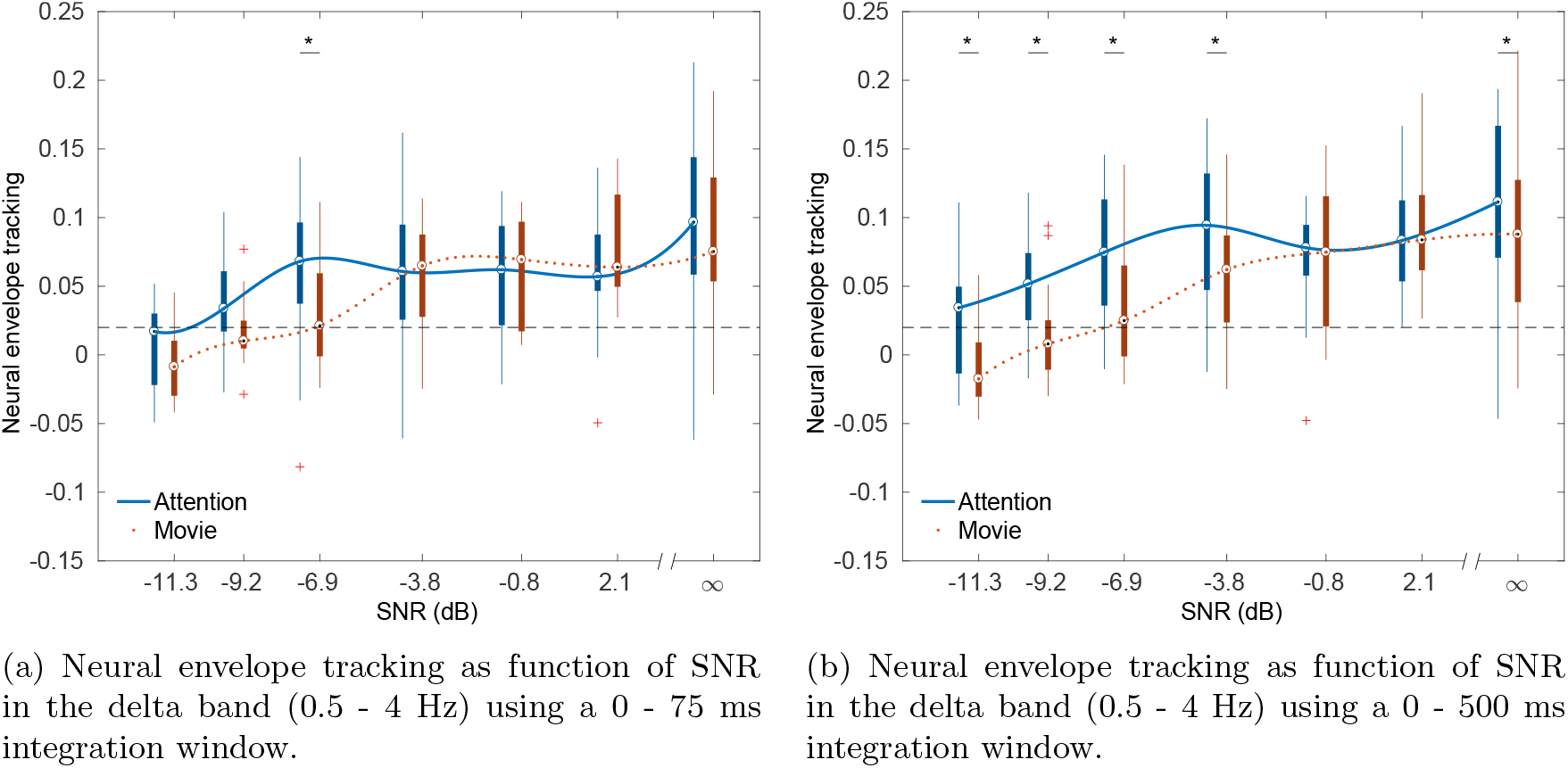
Neural envelope tracking as function of SNR for multiple integration windows.

Using a 0 - 75 ms integration window, we found a significant difference in neural envelope tracking between the attention and movie condition at −6.9 dB SNR using a paired permutation test (*p* = 0.007 after Holm-Bonferroni correction). Using a 0 - 500 ms integration window, we found a significant difference using a paired permutation test between the attention and movie condition at −11.3 dB SNR, −9.2 dB SNR, −6.9 dB SNR, −3.8 dB SNR and in speech in quiet (*p* = 0.007, *p* = 0.024, *p* = 0.024, *p* = 0.024 and *p* = 0.024 after Holm-Bonferroni correction). As we hypothesised that the movie condition would have lower variance than the attention condition, we compared the variance of the neural envelope tracking per SNR. However, while such a trend seens to be present when visually assessing the size of the whiskers in the box plots, using a Brown-Forsythe test, we found no significant differences in spread between attention and movie.

As the attention and movie condition show the expected increase of neural envelope tracking with SNR, we conducted a similar analysis as the behavioural data and attempted to estimate the SRT by fitting a psychometric function on the envelope tracking versus SNR data across subjects. We used the same psychometric curve as for the behavioural data with the exception that the guess-rate and lapse-rate were not fixed. Across subjects, for the attention condition we found an SRT of −9.15 dB SNR (95% confidence interval [−9.34; −8.96] dB SNR). For the watching condition we found an SRT of −6.96 dB SNR (95% confidence interval [−8.57; − 5.35] dB SNR). Compare to the behaviourally measured SRT of −8.03 dB SNR.

#### 3.2.2 Difference in topographies

Figure 2 shows the difference in TRFs between the attention and movie condition in speech in quiet for 17 subjects. After a cluster-based analysis (Maris and Oostenveld, 2007), no significant clusters were found for the story data. The Matrix data however shows a significant difference between the attended and movie condition. The difference from 141 ms to 188 ms is most pronounced in the right-frontal region. The actual TRFs for this interval are shown in Figure 3 and Figure 4. The TRFs are the average of the subject-specific TRFs.

**Figure 2:**
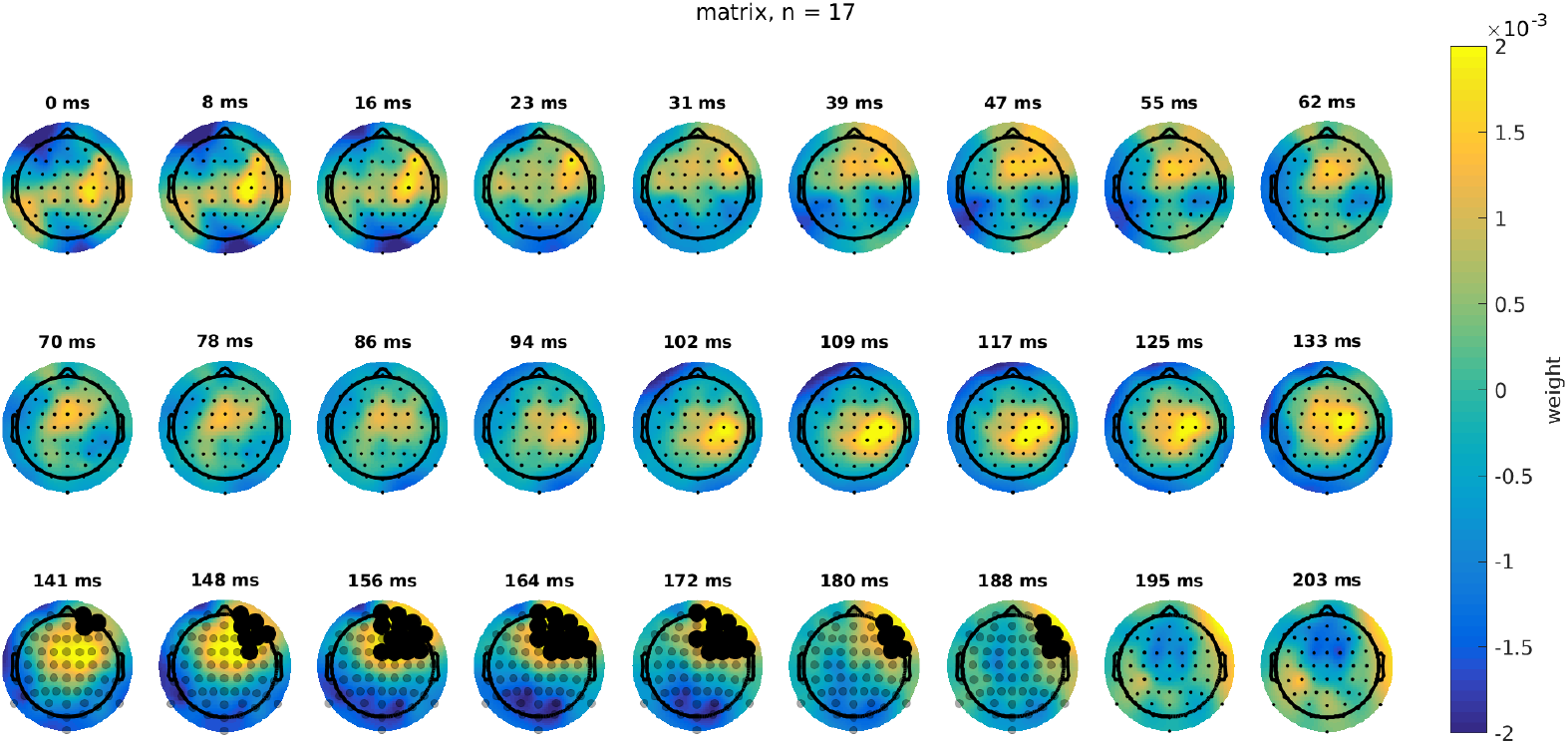
The difference of the topography of the attention condition and the movie condition for the matrix data with a 0 - 500 ms integration window and unfiltered data. The channels part of significant cluster are indicated with black dots.

**Figure 3:**
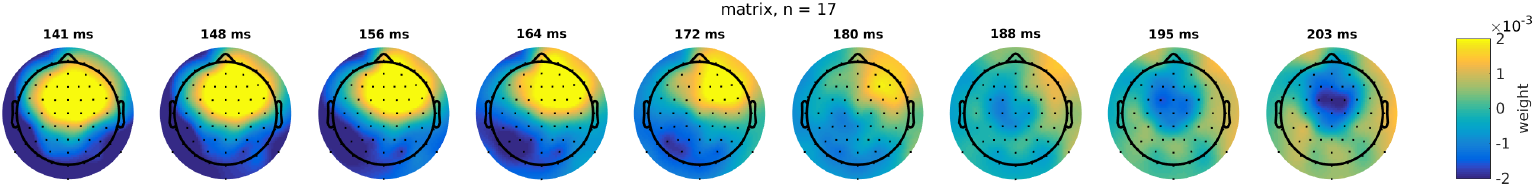
Topography of the attention condition using the Matrix data between 141 - 203 ms.

**Figure 4:**
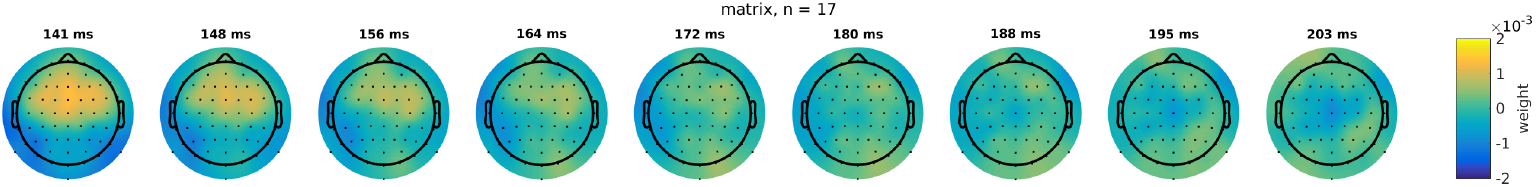
Topography of the movie condition using the Matrix data between 141 - 203 ms.

#### 3.2.3 Difference between TRFs

To better understand why the topographies are different, we averaged the channels contributing most to the difference and did a cluster-based analysis on this TRF. Figure 5 shows the TRF for the attention and the movie condition for the Matrix sentences using a 0 - 500 ms integration window and unfiltered data. 9 participants listened attentively to a story, while in another condition they listened passively to a story while watching a movie. 17 participants did the same for the Matrix sentences. We did a cluster-based analysis to find differences between the TRFs of the two attention conditions (Maris and Oostenveld, 2007). We did not find a significant difference for the story but for the Matrix sentences, a cluster from 141 - 188 ms showed a significant difference. The attended TRF shows a P1 peak at 50 ms, N1 peak at 80 ms and a P2 peak at 160 ms. The movie TRF is similar with the exception that the P2 peak is less pronounced.

**Figure 5:**
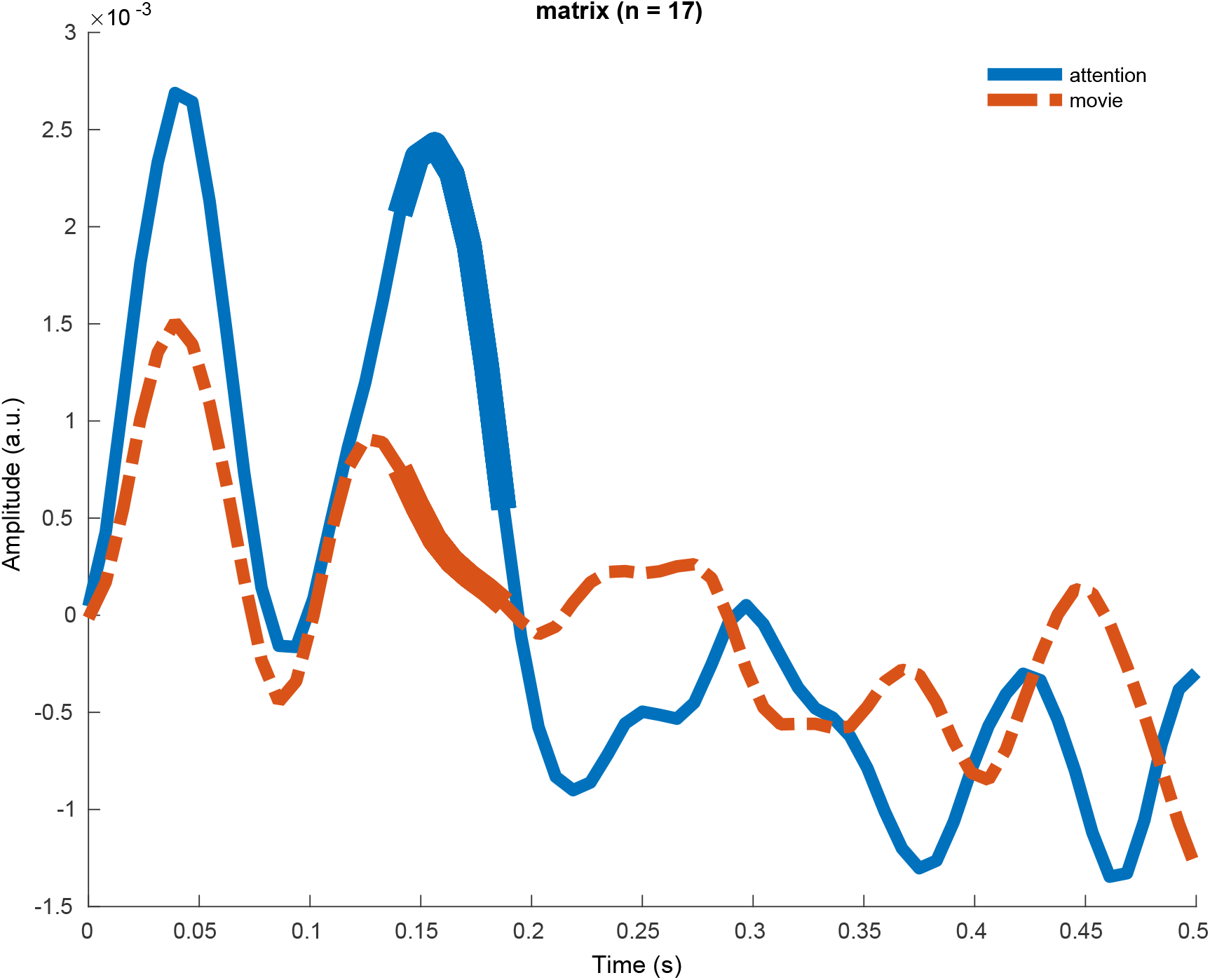
The TRF of the attention and the movie condition for the Matrix sentences using a 0 - 500 ms integration window and unfiltered data. The TRFs are significantly different, the difference is most pronounced from 141 - 188 ms.

### 3.3 Tetris

To further distract the subjects, and direct their full attention to a non-listening task, we also included a condition in which the participants played a computer-game. They were instructed to conduct a visuospatial task, namely play the computer game Tetris, with the difficulty level set such that they were just able to play the game, and had to allocate all of their mental resources to do the task, i.e. the game required maximal effort. They controlled the game using a numeric computer keyboard. This condition was similar to the difficult visuospatial task in Roth et al. (2013a).

We do not present the data because while we found significant neural envelope tracking in the Tetris condition, we did not find a significant correlation between neural tracking and the SNR. This may be because the resources of the participant were exhausted or because of motion artefacts introduced by playing Tetris reducing the EEG signal quality.

## 4 Discussion

We investigated the effect of attention on neural envelope tracking of a single talker, in conditions of maximal attention and watching a silent cartoon movie. To gain insight in the neural origin of the differences, we estimated TRFs and investigated their time course and topography.

### 4.1 Effect of task and SNR

At low SNRs, and therefore lower levels of speech intelligibility, we found higher levels of neural envelope tracking in the attention condition than in the movie condition. This is consistent with the notion that attention requires additional neural resources. The difference in envelope tracking between attention and movie was not visible at higher SNRs, possibly because participants could easily resort to passive listening in these conditions. At low SNRs, passive listening was not possible and the extensive resources needed for active listening were not available due to engagement with the movie. In contrast with our results, Kong et al. (2014) found no significant difference between active and passive listening. This can be explained by the number of participants, the stimulus type and the chosen SNRs. Kong et al. (2014) used 8 participants and a story stimulus, while we have 17 participants for the Matrix condition. The higher amount of subjects gives us more statistical power and Matrix sentences have a very rigid structure compared to a story which may also reduce variance across participants. Also, we measured the attentional effect at more SNRs than Kong et al. (2014). We found that at some SNRs levels the attentional effect is less prominent. Similar to their results, we did not found a significant difference using the story where we used only 9 subjects.

Apart from the level of neural envelope tracking, its relation with SNR and thus speech intelligibility is also important. We saw a consistent increase in neural envelope tracking with SNR for both the attended condition and the movie condition. Unexpectedly, in the movie condition the correlation between SNR and neural level tracking was higher than in the attended condition. We hypothesize that while attention yields higher levels of envelope tracking, it is hard for the subjects to remain fully concentrated during the entire experiment. Therefore, while the movie condition yielded lower envelope tracking levels, it might also have yielded more stable attention, leading to less variability between SNR conditions and therefore a higher correlation between SNR and envelope tracking.

### 4.2 Neural sources related with attention

The literature shows that attentional modulation originates from the parietal cortex and prefrontal cortex (Yantis, 2008; Lavie, 2010a). To investigate this, we assessed the spatial maps of the TRFs for both the attention and movie condition. We found that the difference between the attended TRF and the movie TRF was most pronounced in the right frontal region, which is supported by other research (Alho, 2007).

### 4.3 Latency

In the literature, using the competing talker paradigm, responses earlier than 75 ms are minimally affected by attention while later responses much more so (Snyder et al., 2006; Ross et al., 2009; Choi et al., 2014; Das et al., 2016; Puvvada and Simon, 2017). Using the forward model, we found a significant difference between the attention and movie TRF. This difference was most pronounced from 141 ms to 188 ms. This is in line with the competing talker literature. Moreover, the attended TRF morphology is similar to LAEP literature, showing a P1 peak at 50 ms, N1 peak at 80 ms and a P2 peak at 160 ms. The movie TRF is similar with the exception that the P2 peak is less pronounced. Although the TRFs in quiet at earlier latencies (< 75 ms) does not significantly differ between attention and movie TRF, we still find a significant difference in correlation at some low SNRs.

### 4.4 Distractors

When studying the effect of attention on neural envelope tracking, the difficulty is that there is no reference of the level of attention. Therefore one needs to resort to on the one hand ways to motivate the subject to focus solely on the task at hand (attentively listening to sentences), or on the other hand distract the participants from the task in a controlled way. The latter can be achieved by instructing the participants to ignore the stimulus and providing another task to distract them.

There is a body of research on attention and distraction in auditory and visual tasks. However, most of the literature is focused on how various secondary tasks can distract from a primary task (Murphy et al., 2017; Lavie, 2010b, 2005).

We chose one distractor task: a subtitled cartoon movie. The benefit is that it is engaging and relaxing for the participant. A potential downside is that reading the subtitles requires language processing, which is a resource also required for processing the auditory stimulus. Subtitles are a distractor but maybe they only have a small perceptual load. Another downside is that a movie does not exhaust the available attentional resources of the participants. However, this is difficult to enforce and to quantify how well the participants did.

Murphy et al. (2017) reviewed the literature regarding perceptual load in the auditory domain. A key point of perceptual load theory is that it proposes that our perceptual system has limited processing capacity, and that it is beyond our volitional control as to how much of that capacity will be engaged at any given time. Instead, all of the available information is automatically processed until an individual’s perceptual capacity is exhausted. While this theory is confirmed by many experiments in the purely visual domain, it is less clear in the auditory domain. It seems that the auditory system, with its complicated auditory stream segregation (Shinn-Cunningham, 2008) allows for a less strong selection mechanism. While generally unattended information receives less or no processing, this is not consistently the case. The envelope tracking results with the competing talker paradigm (cf. supra) confirm this and this is also consistent with our results. In the movie TRF, we see that the auditory stimulus is still processed by the brain. However, later processing (from 140 ms on) is reduced.

A concern with using concurrent auditory stimuli to distract from the stimulus of interest is that auditory stream segregation might fail, especially if the listener has an unknown hearing deficit. Therefore a distractor in a non-auditory domain may be desirable. There is some evidence that perceptual load theory also holds across modalities. Macdonald and Lavie (2011); Raveh and Lavie (2015) demonstrated inattentional deafness (failure to notice an auditory stimulus) under visual load, thereby extending the load theory of attention across the auditory and visual modalities, making it clear that vision and audition share a common processing resource, which is consistent with our results. In the movie condition, we saw no P2 peak in the TRF. Also at low SNRs (< −6.6 dB SNR) we found higher neural envelope tracking in the attention condtion compared with the movie condition.

### 4.5 Implications for applied research - Neural envelope tracking as a measure of speech understanding

The stimulus reconstruction method is promising to obtain an objective measure of speech understanding for applications in diagnostics of hearing. Ding and Simon (2013) found a significant correlation between intelligibility and reconstruction accuracy at one signal stimulus SNR. Vanthornhout et al. (2018); Lesenfants et al. (2019) developed a clinically applicable method to objectively estimate the SRT based on reconstruction accuracy, and found a significant correlation between predicted and actual speech reception threshold. In the current study, the average SRT using a behavioural test was −8.0 dB SNR, using the stimulus reconstruction method we found an SRT of −9.2 dB SNR in the attention condition and an SRT of −7.0 dB SNR in the movie condition. Both objective methods are thus close to the behavioural method. The rank order of these SRT values also makes intuitive sense: when paying maximal attention, and estimating the SRT based on envelope tracking, the lowest value (best performance) is obtained. In the behavioural experiment, the subjects’ neural resources are taxed more, because they need to repeat each sentence, leading to a slightly higher SRT. Finally, when not paying attention to the stimulus, the SRT reaches its highest value.

We found that neural envelope tracking was better correlated with SNR in the movie condition, which is important for the estimation of the SRT. While the level of attention influenced the level of neural tracking, to obtain an objective measure of speech understanding it may be more important to have stable attention and thus lower variability, than to have high levels of neural envelope tracking. Equalising the level of attention across subjects and conditions by giving them an unrelated and/or easy task, such as watching a movie might therefore yield the best results.

In addition to procedural benefits, an additional benefit of letting subjects watch a movie instead of attending to low-content sentences, is that it is much more pleasant for the subjects, and therefore easier to implement in the clinic, especially in populations such as children.

## 5 Conclusion

Using a movie, we varied the level of attention while we estimated the neural envelope tracking. Neural envelope tracking was significantly different between the condition with maximal attention and the movie condition. This difference was most pronounced in the right-frontal region of the brain. However, this does not seem to be a problem for estimating speech understanding as neural envelope tracking was still highly correlated with the stimulus SNR, even more in the movie condition.

## 6 Acknowledgements

The authors thank Lise Goris, Lauren Commers, Paulien Vangompel and Lies Van Dorpe for their help with the data acquisition. Financial support was provided by the KU Leuven Special Research Fund under grant OT/14/119 to Tom Francart. Research funded by a PhD grant of the Research Foundation Flanders (FWO). This project has received funding from the European Research Council (ERC) under the European Union’s Horizon 2020 research and innovation programme (grant agreement No 637424), from the YouReCa Junior Mobility Programme under grant JUMO/15/034 to Jonas Vanthornhout. The authors declare that they have no conflict of interest.

## Footnotes

1 The maximal deviation between the shown SNR and the actual SNR was 2.5 dB.

